# Creating overlapping genes by alternate-frame insertion

**DOI:** 10.1101/2024.11.07.622342

**Authors:** Sean P. Leonard, Tiffany Halvorsen, Bentley Lim, Dan M. Park, Yongqin Jiao, Mimi Yung, Dante Ricci

**Affiliations:** Biosciences and Biotechnology Division, Physical and Life Sciences Directorate, Lawrence Livermore National Laboratory, Livermore, CA 94550, USA

## Abstract

Overlapping genes–wherein two different proteins are translated from alternative frames of the same DNA sequence–provide a means to stabilize an engineered gene by directly linking its evolutionary fate with that of an overlapped gene. However, creating overlapping gene pairs is challenging as it requires redesign of both protein products to accommodate overlap constraints. Here, we present a new “<u>o</u>verlapping, <u>a</u>lternate-<u>f</u>rame insertion” (OAFI) method for creating overlapping genes by *insertion* of an “inner” gene, encoded in an alternate frame, into a flexible region of an “outer” gene. Using OAFI, we create new overlapping gene pairs of bacterial toxins within an antibiotic resistance gene. We show that both the inner and outer genes retain functionality despite redesign, with translation of the inner gene is influenced by its overlap position in the outer gene. Additionally, we show that selection for the outer gene alters the permitted inactivating mutations in the inner gene and that overlapping toxins can restrict horizontal gene transfer of the antibiotic resistance gene. Overall, OAFI offers a versatile tool for synthetic biology, expanding the applications of overlapping genes in gene stabilization and biocontainment.

## Introduction

Since overlapping genes were first discovered in the genome of phage φX174 in the 1970’s^1,2^, they have represented a unique means of compressing genetic information, regulating gene expression, and tuning molecular evolution^3^. Defying the historical “one gene-one enzyme” dogma^4^, these stretches of DNA encode multiple, overlapping coding sequences in alternate reading frames, can be situated on the sense or antisense strand of DNA, and can be found across the tree of life, particularly in viral genomes^5^. Mutations within overlap regions impact the sequence— and, therefore, function— of both genes simultaneously. Often, mutations that are synonymous in one frame will be non-synonymous in the alternate frame. Naturally occurring gene overlaps narrow the fitness landscape each gene can explore and can ultimately increase mutational robustness^6^. In RNA viruses, which are subject to naturally higher mutation rates than DNA viruses and frequently encode overlapping genes, this narrowed mutational landscape reduces the frequency of synonymous mutations and increases fitness^6,7^.

The unique evolutionary and compositional constraints of natural overlapped genes are reasonably well understood^8–10^, enabling several groups to predict the potential applied utility of artificial, use-inspired gene overlaps^5,11^. Synthetic biology has rapidly advanced in recent years, but major challenges remain in creating *genetically stable* designs that can reliably perform in complex environments without mutational loss of function or unintended dissemination of genetically engineered DNA (i.e., horizontal gene transfer or HGT)^12^. Gene overlaps may help solve these problems, as artificially created overlaps have the potential to alter the evolutionary fate of a target gene, a useful property to stabilize and contain synthetic genetic devices.

Creating overlapping genes *de novo* remains a daunting challenge, however, despite novel computational tools for designing overlapping genes^13,14^. This is largely a product of the requirement to extensively redesign one or both entangled ORFs at the amino acid level when generating gene overlaps^13^. A comparison of theoretical, designed overlapping sequences to their naturally-occurring homologs showed that the sequences of variants designed to enable overlap are generally similar to, and sometimes indistinguishable from, natural homologs within a gene family^15^, suggesting their *de novo* design is possible but at the limits of current protein design methods. Moreover, designed overlapping genes often are not functional when tested in the laboratory. For example, Wang and coworkers successfully created new overlapping genes, but after screening > 7500 predicted designs identified just 6 functional designs, a success rate of < 0.1%^13^. Factors underlying failed overlap designs likely include poor sequence conservation, incorrect protein folding, codon usage, and overlap position, all of which are understudied. Successful design of overlapping genes thus remains difficult and error-prone, and the lack of reliable methods to create them hinders our ability to study and develop them for biotechnological applications.

To enable on-demand access to synthetic gene overlaps, we hypothesized that we could directly create overlapping genes, with minimal amino acid changes to both genes, through *insertion* of one open reading frame into another encoded in an alternate frame. Here we demonstrate the creation and evaluation of such “<u>o</u>verlapping, <u>a</u>lternate-<u>f</u>rame insertions” (“OAFI”), show that function of both genes can be retained despite this redesign, and show that selection on the overlapping partner alters the permitted inactivating mutations of a costly gene. Our strategy thus represents a new, facile method for creating artificial gene overlaps to shape the evolutionary trajectory of synthetic devices, one that may be readily extended to various biotechnology applications.

## Results

### Design of overlapping, alternate-frame insertions

Our approach to generate OAFIs requires (1) identifying putative insertion-tolerant sites of an “outer gene”, (2) recoding a desired “inner gene” to remove alternate-frame stop codons, and then (3) inserting the inner gene within an alternate reading frame of the outer gene (**Figure 1A**). For our initial designs we selected the tetracycline resistance factor *tetA* from Tn10^16^ (**Figure 1B**). *tetA* encodes a multi-pass inner membrane efflux pump that promotes drug resistance in Gram-negative bacteria by selectively exporting tetracycline from the cytosol. As some loops connecting transmembrane domains can accommodate considerable variation in length^17,18^, we reasoned these loops could be viable OAFI targets if superfluous amino acids can be added to the *tetA* frame at a loop without disrupting TetA function. Based on the Alphafold^19^ structure available at UniProt (AF-P02980-F1) and prior topological analysis of TetA^18^, we targeted three locations in non-membrane-spanning regions of TetA for gene insertions: an amino-terminal periplasmic site (“early”, after S36), a large cytosolic loop near the midpoint of the protein sequence (“middle”, after G195), and one carboxy-terminal cytosolic site (“late”, after Q391) (**Figure 1C**). The middle and late positions are associated with low pLDDT in the predicted structures (< 50), possibly suggesting more structural flexibility and therefore more suitability for insertion. The insertions do not remove any amino acids present in the original TetA sequence but do introduce new amino acids according to the length of the inserted gene. We also designed an “operon control” which encodes each inserted gene immediately after the stop codon of *tetA*. The operon control should retain the transcriptional and translational profiles of the overlapped genes but without overlapping coding sequence, thus controlling for the impact of *overlap* as opposed to genetic linkage (i.e., proximity). Throughout this work, the *tetA* gene is driven by its native promoter and ribosome binding site (RBS), and the inner genes (including the operon control) all feature the same strong 18-bp RBS upstream of the inner ORF. Inner genes were recoded to remove stop codons in the native +2 frame (in-frame with *tetA*) and then inserted into the *tetA* gene in the +3 frame.

**Figure 1.**
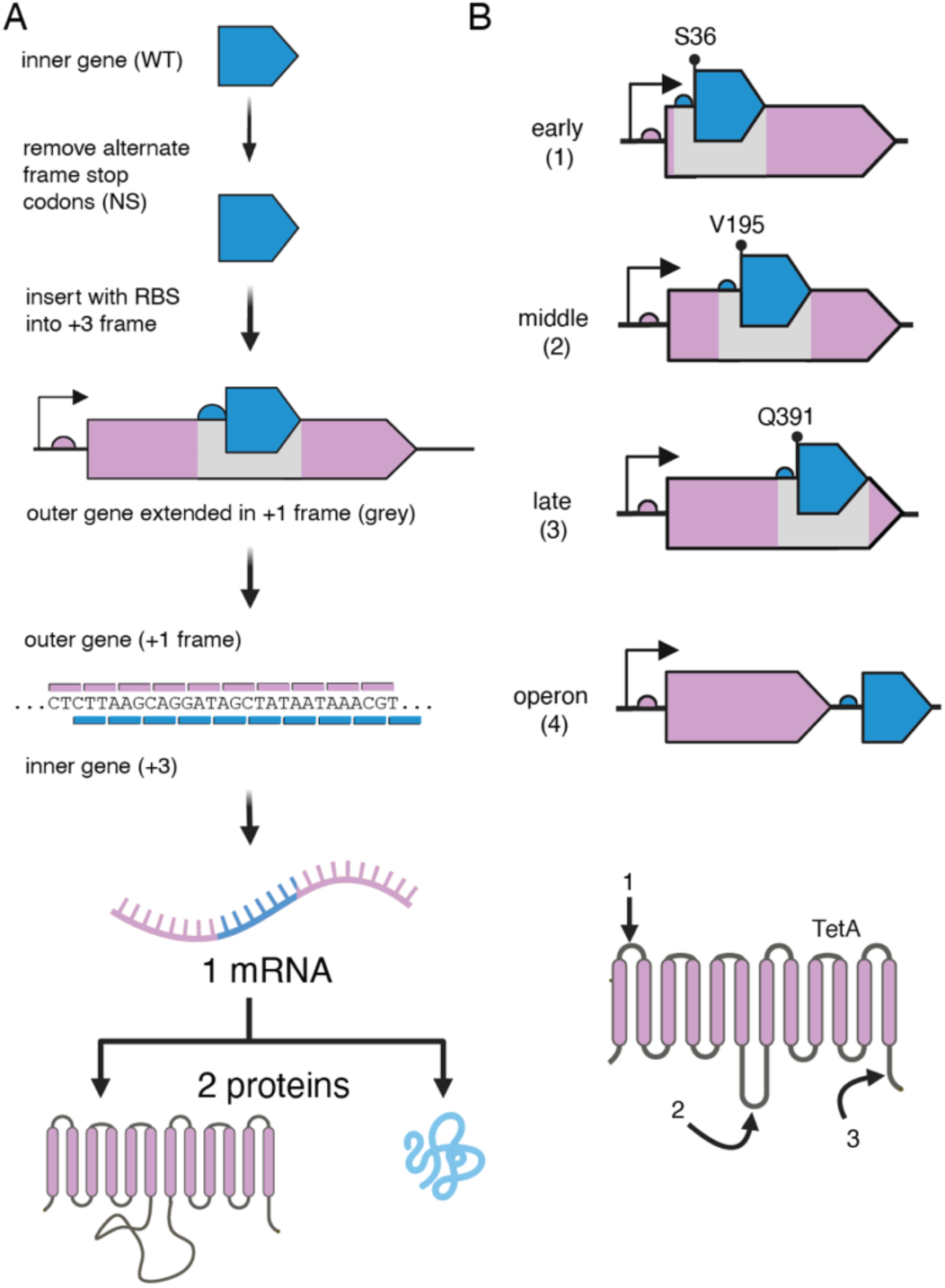
Creating overlapping genes with alternate frame insertions. (A) After identifying the desired inner (blue) and outer (lilac) gene pair, the inner gene is recoded to remove stop codons in the native +2 frame before its insertion into a permissive location of the outer gene in the alternate frame. An RBS is included in the inner gene’s reading frame to enable separate translation of both frames. (B) Example gene pair used in this study, with inner genes inserted into 3 positions of *tetA* at (1) early, (2) middle, (3) late, and (4) a downstream operon position. Arrows (bottom) correspond to approximate insert locations on TetA topology.

### Overlapping toxins limit recovery of transformed cells

We created our initial overlapping, alternate-frame insertions to prevent the HGT-mediated dissemination of an antibiotic resistance gene by “poisoning” it with an overlapping bacterial toxin. We selected toxins *hicA* and *tse2* as inner genes because the gene products vary in length (59 and 159 amino acids (aa), respectively) and in mechanism of action, and they require different threshold concentrations to arrest growth^20,21^. We made synonymous substitutions in 4 codons each of *hicA* and *tse2* to yield the “NS” (no stops) alleles for each toxin, which lack stop codons in the native +2 frame. To successfully clone these toxins, we created “immune” strains constitutively expressing the corresponding antitoxin from the *att*Tn7 site in NEB5alpha. To confirm the functionality of the recoded alleles, we placed them under control of the acyl homoserine lactone (AHL, 3-oxo-C6-HSL) inducible *P_lux_* promoter and measured toxicity by transformation into toxin-sensitive hosts with or without induction of the toxin^11^(**Figure 2A**). We quantified colony forming units (CFU) following toxin plasmid transformation compared to control transformations using a known non-toxic construct (*gfp*) and observed a ∼10,000-fold reduction in transformation efficiency associated with the toxin genes, with few or no resulting CFUs recovered. No significant differences were observed for transformations using *hicA*^NS^ and *tse2*^NS^ constructs relative to those using parental genes (*hicA*^WT^ and *tse2*^WT^). *hicA*^WT^ and *hicA*^NS^ transformants were recovered when the AHL inducer was omitted from the growth medium, but the *tse2*^WT^ and *tse2*^NS^ constructs were lethal even without induction, consistent with the previously reported potency of the Tse2 toxin^21^.

**Figure 2.**
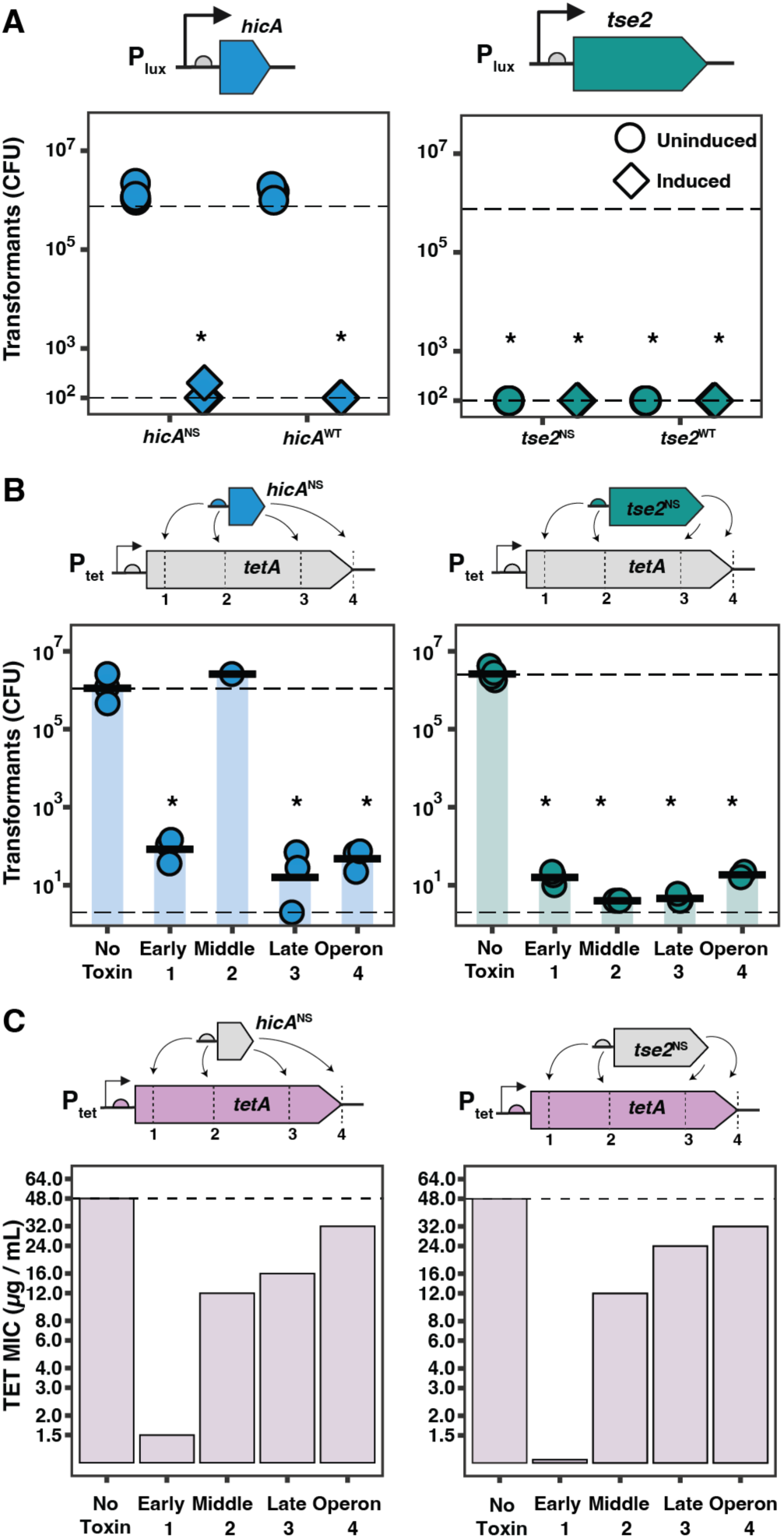
Alternate-frame insertion creates functional overlapping gene pairs. (A) *hicA* (left) and *tse2* (right) variants recoded to remove alternate frame stop codons (“no stops”, NS) retain function by transformation toxicity assay. Upper dotted line corresponds to number of transformants obtained with non-toxic plasmid encoding *sfgfp*, lower dotted line corresponds to approximate limit of detection. Tse2 shows the greatest toxicity, as even uninduced transformants do not grow. (B) Transformation assays of overlapping constructs with *hicA*^NS^ (left) or *tse2*^NS^ (right) inserted into TetA. HicA inserts are toxic in all positions except the middle position, while Tse2 remains toxic at all inserted locations. Upper dotted line corresponds to number of transformants obtained with plasmid bearing wild-type TetA with no toxin. Lower dotted line corresponds to approximate limit of detection. (C) TET MIC for wild-type allele and overlapping designs for HicA (left) and Tse2 (right). Asterisk indicates statistically significant differences between groups (ANOVA followed by Tukey HSD, 𝑝 < 0.05).

We next inserted *hicA*^NS^ and *tse2*^NS^ into *tetA* at the three internal test positions and downstream operon control and then assessed both overlapping gene partners for each functional gene product, i.e., tetracycline (TET) resistance (through minimal inhibitory concentration (MIC) assays) and bacterial toxicity (through transformation toxicity assays) (**Figure 2B, C)**. The operon constructs exhibited slightly lower TET MIC (32 μg/mL) than WT *tetA* (48 μg/mL).

Insertion of either toxin at any of the target insertion sites in *tetA* led to a reduction in TetA activity, with the early insertions being most disruptive to TetA function (MIC of 1.5 μg/mL for *hicA*^NS^, 0.5 μg/mL *tse2*^NS^) (**Figure 2C**). Middle and late position insertions maintained a TET MIC of >10 μg/mL, which is suitable for selection in *E. coli*. Despite having markedly different lengths and sequences, the *tse2*^NS^ and *hicA*^NS^ insertions have similar impacts on TetA function. In fact, in one instance a longer (*tse2*^NS^) insertion seemed to compromise TetA function less than the shorter (*hicA*^NS^) insertion (late position, 24 μg/mL vs. 16 μg/mL respectively).

To assess toxin function, we transformed plasmids encoding the overlapping pairs into toxin-susceptible *E. coli* cells and compared the transformation frequencies to those observed using a plasmid encoding only *tetA* (**Figure 2B**). Since the upstream RBS of each inserted inner gene is identical, we initially expected toxin expression to be similar across overlap positions. As expected, *tse2*^NS^ was toxic at all insertion locations in *tetA*, comparable to the operon design, indicating that *tse2*^NS^ is actively expressed and sufficiently potent to prevent growth. However, *hicA*^NS^ was toxic only when inserted into the early and late positions of *tetA* and appeared non-functional in the middle position. As toxicity seems to require more *hicA*^NS^ expression relative to *tse2*^NS^ (**Figure 1A**), we hypothesized that insertion at the middle position may lead to reduced expression of functional toxin.

### Overlap position impacts inner gene translation

The observation of an unexpectedly non-functional insertion of *hicA*^NS^ led us to explicitly investigate the positional effects of alternate frame insertion on translation of the inner gene. We created an alternate-frame translational reporter in *tetA* that encodes the 238-aa superfolder GFP fluorescent protein (altered through 10 silent mutations and one missense mutation, L78M, to eliminate out-of-frame stop codons), and we measured the impact of overlap design on function of both the inner gene (GFP/fluorescence; **Figure 3A**) and the outer gene (TetA/MIC; **Figure 3B**). Placement of the recoded reporter gene (*gfp*^NS^) downstream of *tetA* did not alter the TET MIC (48 μg/mL). As with the toxin overlaps, *gfp*^NS^ insertion at any position within *tetA* reduced TET MIC to varying degrees. The most dramatic decrease in MIC occurred with *gfp*^NS^ inserted in the early position, which completely disrupted TetA function (MIC < 0.5 μg/mL). GFP insertion at the middle and late positions resulted in a more pronounced reduction in MIC than either toxin insert (8 μg/mL middle, 12 μg/mL late), perhaps due to the increased length of *gfp*^NS^ relative to *hicA*^NS^ and *tse2*^NS^. Of note, GFP fluorescence does not appear to inversely correlate with MIC; both early and late insertions have increased fluorescence compared to the operon position, but the late position is substantially more fluorescent and retains a higher MIC than both early and middle positions. The middle insertion has the lowest GFP fluorescence, consistent with the apparent reduction in HicA function when expressed from the same position (**Figure 2B**). Altogether, these data indicate that the functionality of TetA is affected by insertional overlaps, with the late (C-terminal) insertions being the most well-tolerated, and that expression of the inner gene can be modulated based on its position within the outer gene.

**Figure 3.**
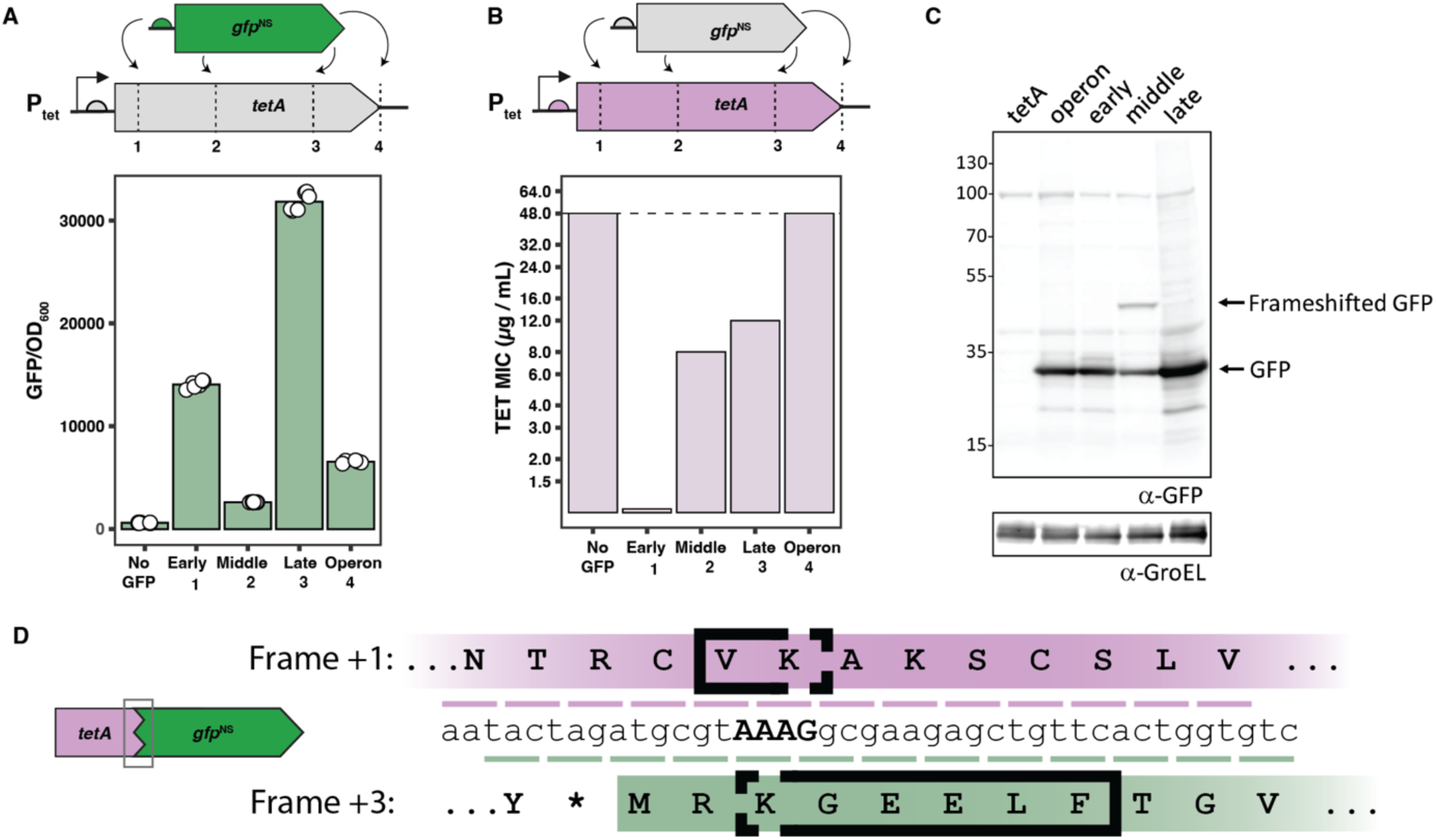
Overlap position alters translation of inner gene. (A) Fluorescence of overlapped *gfp*^NS^ constructs in *tetA*. Expression significantly varies at each position and is highest at the late position. (B) TET MIC for wild-type allele and overlapping *gfp*^NS^ designs. TET MIC is highest for the WT allele and operon controls. Insertion at the early position of *tetA* disrupts TetA function. (C) Western blot of GFP translated from TetA insertions. GFP is translated regardless of position in the outer gene, but the middle position also produces a fusion peptide from a translational frameshift. (D) Schematic of TetA-GFP fusion product with identified fusion peptide (black box). Ribosomal frameshift motif (X-XXY) bolded. All groups in (A) are significantly different from one another (ANOVA followed by Tukey HSD, 𝑝 < 0.05).

To determine if the wide range of fluorescence intensities across *gfp*^NS^ insertions is due to differences in protein abundance, we examined the levels of GFP translated from all three overlap locations by western blotting (**Figure 3C**). All constructs produced GFP at the expected size (∼27 kDa), indicating that overlapping GFP can be translated independently from TetA to yield a single protein product. These relative GFP protein levels varied for each position, and these abundances correlate well with *in vivo* fluorescence (**Figure S1**). We confirmed that these differences were not attributable to transcriptional differences, as transcript abundance was found to be comparable across constructs (**Figure S2**). Intriguingly, we detected a higher molecular weight band using anti-GFP antibodies (∼48 kDa) only when *gfp*^NS^ is translated from the middle position of *tetA* (**Figure 3C**). Based on the size of this band, we hypothesized that this product might comprise a fusion of TetA and GFP that results from a translational frameshift. To test this hypothesis, we used a monoclonal anti-GFP antibody to immunoprecipitate GFP from the strain encoding *gfp*^NS^ in the middle position of *tetA*, specifically isolated the co-precipitating ∼48 kDa product, and analyzed the purified protein variant via tandem mass spectrometry (MS/MS)^22^.

Peptides were detected against the amino acid sequence 1) in the N-terminus of TetA encoded upstream of the *gfp*^NS^ overlap region, and 2) throughout the length of GFP (**Dataset 1**). This finding suggests that the ∼48 kDa band represents a TetA-GFP fusion. Moreover, we identified a unique peptide (VKGEELFT) not found in either TetA or GFP that could only arise from fusion of amino acids from the +1 frame and the +3 frame at the third codon of GFP, specifying the junction between the first 201/2 amino acids in the +1 frame and residues 3/4 of GFP in the +3 frame (**Figure S3**). Because of the shared K residue, we do not know whether K was encoded in the +1 or +3 frame.

Further analysis of the nucleotide sequence of the fusion peptide suggests a -1 frameshift occurring at an A-AAG sequence, just downstream of the RBS for *gfp*^NS^. These sequence features are consistent with frameshift sites in natural -1 programmed ribosome frameshift events commonly found in viruses and bacteria which typically occur at “slippery” X-XXY-YYZ motifs downstream of an RBS^23^. The A-AAG sequence found within *gfp*^NS^ only has a partial X-XXY motif which may explain the lower prevalence of the ∼48 kDa product relative to the GFP product at ∼27 kDa. Overall, these results show the possibility for unexpected, complex frameshifted products to form during construction of overlapping genes and suggest low expression of the inner gene from the middle position. The latter may explain the lack of toxin functionality observed with the middle *hicA*^NS^ insertion.

### Selection for outer gene function limits rates and mechanisms of inner toxin escape

Gene overlaps can link the evolutionary trajectory of an unselected ORF to the selection pressures imposed on an overlapping ORF, an effect we refer to as “overlap epistasis”^6,7^. As our OAFI designs comprise both a selectable marker (*tetA*) and a counterselectable marker (*tse2*^NS^), we sought to determine the extent to which our OAFI designs exhibit overlap epistasis by measuring how selection for the outer gene affects evolution of the inner gene. We focused on the *tse2*^NS^ designs that retained both Tse2 toxin activity and TetA function (i.e., middle, late, and operon constructs). We hypothesized that overlap with the conditionally essential *tetA* would constrain the spectrum of *tse2*^NS^-inactivating mutations when *tetA* function was selected for (i.e., in the presence of TET). As before, we transformed *E. coli* with plasmids encoding *tetA*::*tse2*^NS^ and plated transformants without TET (“–TET”, selecting only for plasmid uptake) or with TET (“+TET”, dually selecting for plasmid uptake and TetA function) and interpret all viable colonies to represent toxin “escape” (e.g., mutational inactivation of the toxin to permit growth). As expected, in the absence of an overlapping toxin, selection for *tetA* (+TET) had no impact on the number of observed transformants (**Figure 4**). When *tse2*^NS^ was present, however, selection for TetA significantly reduced the number of escapees in the middle and operon constructs. This effect was most pronounced for the middle position construct, which yielded no transformants after 24 hours in +TET, a >3 log-fold decrease from –TET. This indicates the relative rarity of mutations that inactivate Tse2 but retain normal TetA function when they are overlapped in this middle arrangement. We did, however, observe several tiny colonies on +TET plates with the middle construct after prolonged incubation (> 72 hours), suggesting that these colonies carry mutations that only partially preserve TetA function and therefore exhibit only partial resistance to tetracycline, resulting in slow growth. Compared to the middle position of *tetA,* differences in escape frequency observed between +TET and –TET are modest for the late and operon constructs (**Figure 4**), indicating that while selection for TetA may preclude certain inactivating mutations (e.g., large deletions covering most or all of *tetA*), null mutations in the toxin that preserve TetA function can still arise at a high frequency in these constructs.

**Figure 4.**
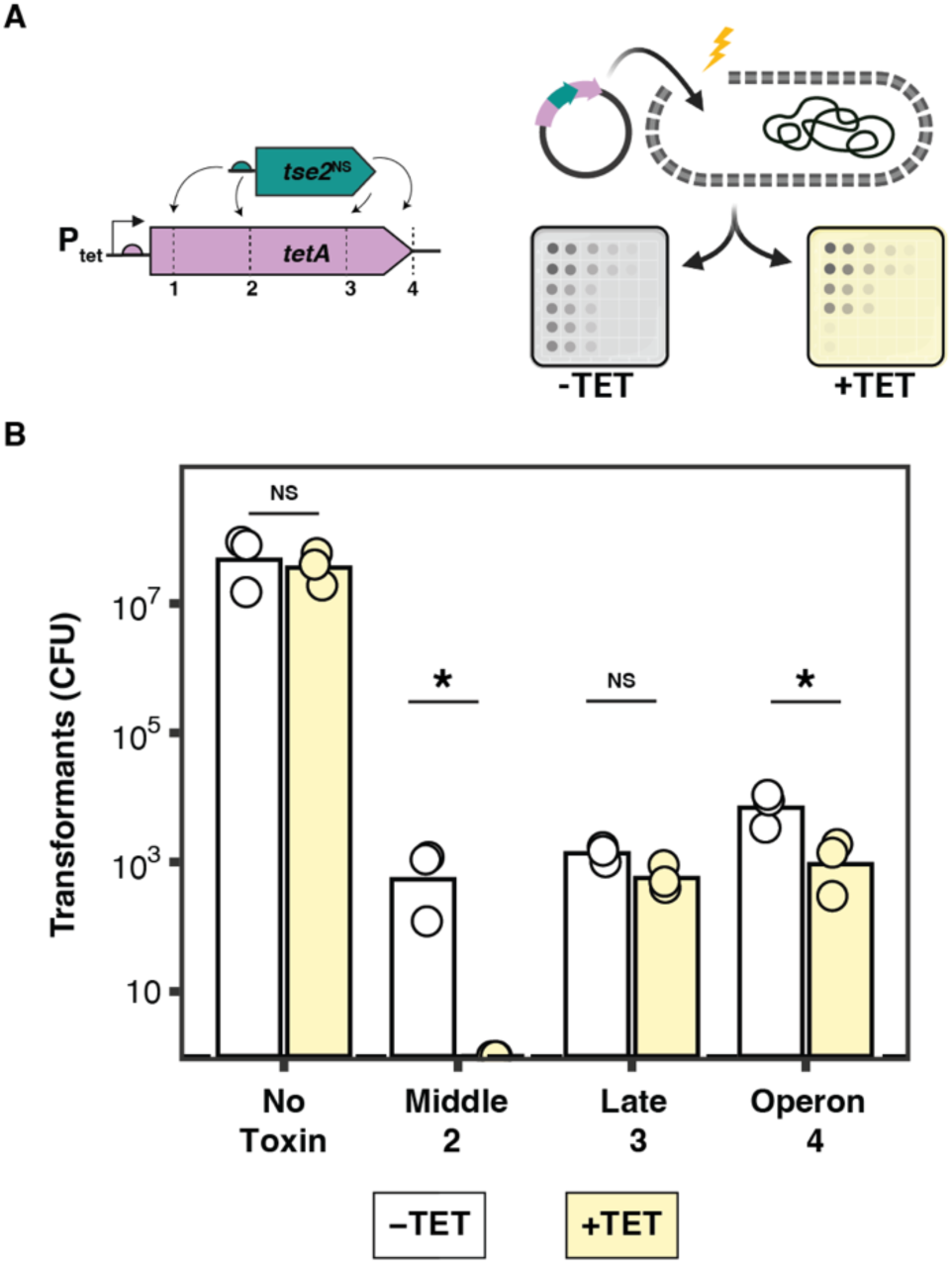
TetA selection decreases escape rate of overlapped toxin Tse2. (A) Transformability and toxin escape of overlapped *tse2*^NS^ in the presence or absence of external gene selection (–TET / +TET). (B) Selection for functional TetA lowers escape rate in overlapped *tse2*^NS^ constructs at the middle insertion location. Asterisks indicated statistical differences between groups (ANOVA followed by Tukey HSD, 𝑝 < 0.05).

To clarify the impact that selection for the outer gene (TetA) had on the spectrum of inactivating mutations for Tse2, we sequenced multiple escaped colonies (Tse^-^) from each transformation with (Tet^R^) and without TET selection (8-13 clones/condition, 96 clones in total). In 83/96 clones (86.4%), we identified a single, presumably causative escape mutation, 73 of which were determined to be unique and independent after accounting for experimental grouping and sampling replicates. The most common mutations were deletions (46/73) and mobile element insertions (21/73). In the –TET condition, we predominantly observed mobile element insertions or large deletions encompassing both *tetA* and *tse2*, regardless of the insertion position (**Figure 5A**). This is consistent with the lack of any selection pressure to maintain tetracycline resistance. In contrast, +TET breakthrough colonies contained mutations in the *tse2*^NS^ insertion, with none encompassing large regions of *tetA*. This difference is most apparent for the middle construct, wherein only small deletions (<= 36 bp) and single-nucleotide polymorphisms or insertions were identified from the small colonies that arose after 72 hours on +TET selection (Figure 5B). These escapees had no large deletions or mobile element insertions and 11/14 identified mutants were in-frame. This marked shift in the spectrum of permitted Tet^R^ Tse^-^ variants of the middle construct likely explains the greatly reduced transformation frequency seen with +TET selection (**Figure 4B**). In the late construct, the deletions found in Tet^R^ Tse^-^ escape mutants often also included the cytosolic C-terminus of TetA, suggesting that this peptide is dispensable for efflux. In the operon construct, several identical mutations were found in the +TET and –TET conditions, indicating minimal impact of TET selection on the observed mutational spectrum.

**Figure 5.**
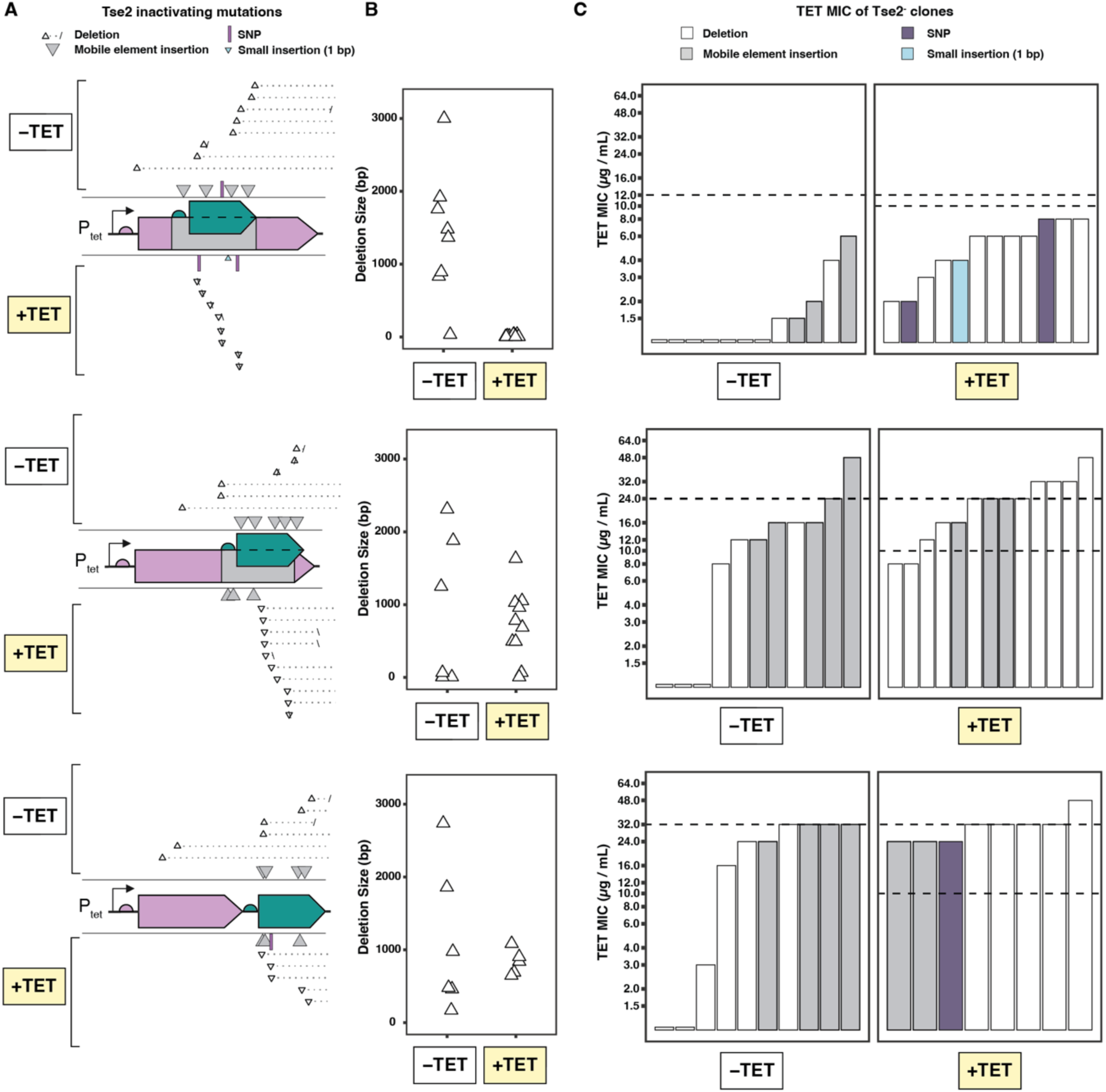
Selection for outer gene alters mutational spectra of toxin escape. (A) Unique mutations identified in individual colonies of escaped transformants from Figure 4 with (+TET) or without (–TET) selection for the outer gene (TetA) in middle (top), late (middle), and operon (bottom) configurations. Unique mutations are plotted at the corresponding approximate location in the overlapping gene construct: deletions (white triangles), mobile element insertions (grey triangles), single nucleotide polymorphisms (purple bars), and small insertions (blue triangle). Deletions are truncated immediately downstream of construct to aid visualization. (B) Size difference in deletions shown in (A). In the middle position, +TET leads to significantly smaller deletions. (C) TET MIC of plasmids bearing mutations shown in (A), measured after retransformation.

To quantitatively assess TetA function in escape variants, we retransformed the purified plasmids from Tse2^-^ colonies into susceptible hosts and measured TET MIC (**Figure 5C**). In the middle construct, mutations isolated from –TET abolished TetA function (7/12 clones), while the remaining reduced MIC to < 6 μg/mL. In contrast, for both the late and operon constructs many Tse2^-^ mutants isolated from –TET remained Tet^R^ (MIC > 10 μg/mL), although for each of the constructs, 2 or 3 mutants were Tet^S^. These results confirm that spontaneous *tse2* inactivating mutations can also inactivate *tetA* function. However, in contrast to the late and operon constructs, in which Tse^-^ mutants are often phenotypically Tet^R^, none of the mutations isolated in the middle construct that inactivated the toxin also preserved normal TetA function. Even when selecting for tetracycline resistance, we find that all Tse^-^ mutants for the middle construct had MIC between 2-8 μg/mL, which is consistent with them requiring > 72 h to grow on +TET plates. However, these MIC values were higher than those observed for the –TET condition, supporting that TET selective pressure retained at least partial TetA function in all escapees.

Nearly all mutations identified in the +TET condition for the late and operon constructs retained MIC > 10 μg/mL, indicating that many mutations for these constructs inactivate toxin function but retain TetA function. These included several large mobile element insertions in *tetA* and large deletions that deleted the C-terminus of TetA, confirming that alteration or removal of the TetA C-terminal tail has minimal impact on TET MIC, as has been suggested previously^24^.

To understand the impact that selection strength has on mutational robustness of the inner gene, we performed an additional set of experiments in which we selected with a lower concentration of tetracycline (5 μg/mL) (**Figure S4**). These results resemble those presented in **Figure 5**, in that (1) toxin-inactivated mutations are dominated by large deletions and mobile elements and (2) SNPs were observed only in the middle position in the +TET condition, indicative of Tet^R^ Tse^-^ mutations, and (3) mutants isolated on +TET maintained growth on tetracycline. However, decreased stringency of selection for TetA function does allow some mobile element insertion mutations to occur in the middle position in the +TET condition. Overall, these results demonstrate that by increasing selection pressures on the outer gene, alternate frame insertions can greatly reduce mutational inactivation of a gene of interest by restricting the diversity of permitted inactivating mutations in a position-dependent manner.

### Overlapping toxins limit horizontal gene transfer of antibiotic resistance genes by natural transformation

Next, we sought to determine whether our overlapping *tetA*-*tse2*^NS^ designs reduce rates of natural transformation—an important route for HGT in natural populations^25,26^. Unlike electroporation, natural transformation occurs broadly in nature and often proceeds through single-stranded intermediates that may alter the frequency of escape mutations^27–29^. We transferred the middle *tetA*::*tse2* ^NS^ overlapped design to a broad-host-range plasmid^30,31^ and tested its transfer into naturally competent strains of the soil bacterium *Acinetobacter baylyi* using an agar plate-based natural transformation protocol. We selected *A. baylyi* ADP1 for its high rates of nonspecific DNA uptake^32^, along with its derivative ADP1-ISx, which lacks active mobile elements^33^. As expected, both ADP1 and ISx were readily transformed by a control plasmid harboring wild-type *tetA* at high rates (0.9 × 10^4^ CFU/μg and 1.8 × 10^4^ CFU/μg, respectively), independent of tetracycline selection (**Figure 6**). As observed in *E. coli*, the transformation rate of the *tetA-tse2*^NS^ middle construct into either *A. baylyi* strain was significantly reduced (∼100-fold) compared to the *tetA*-only control owing to the presence of the lethal *tse2*^NS^ gene, with no colonies recovered after 24 hours in the +TET condition. After 48 hours, we observed numerous additional small colonies growing in the –TET condition, suggesting that Tse2 may be less toxic to *A. baylyi* than to *E. coli*; these strains were not characterized further. No such colonies arose in the +TET condition. We observed no difference in the transformation frequency of the *tetA-tse2*^NS^ middle construct between ADP1-ISx and wild-type ADP1 in either TET condition, suggesting that mobile element insertion is not the major driver of mutational escape from Tse2 toxicity. Altogether, these findings indicate that alternate-frame insertion is a promising approach for designing synthetic DNA constructs that resist HGT in environmental microbes and natural microbial communities.

**Figure 6.**
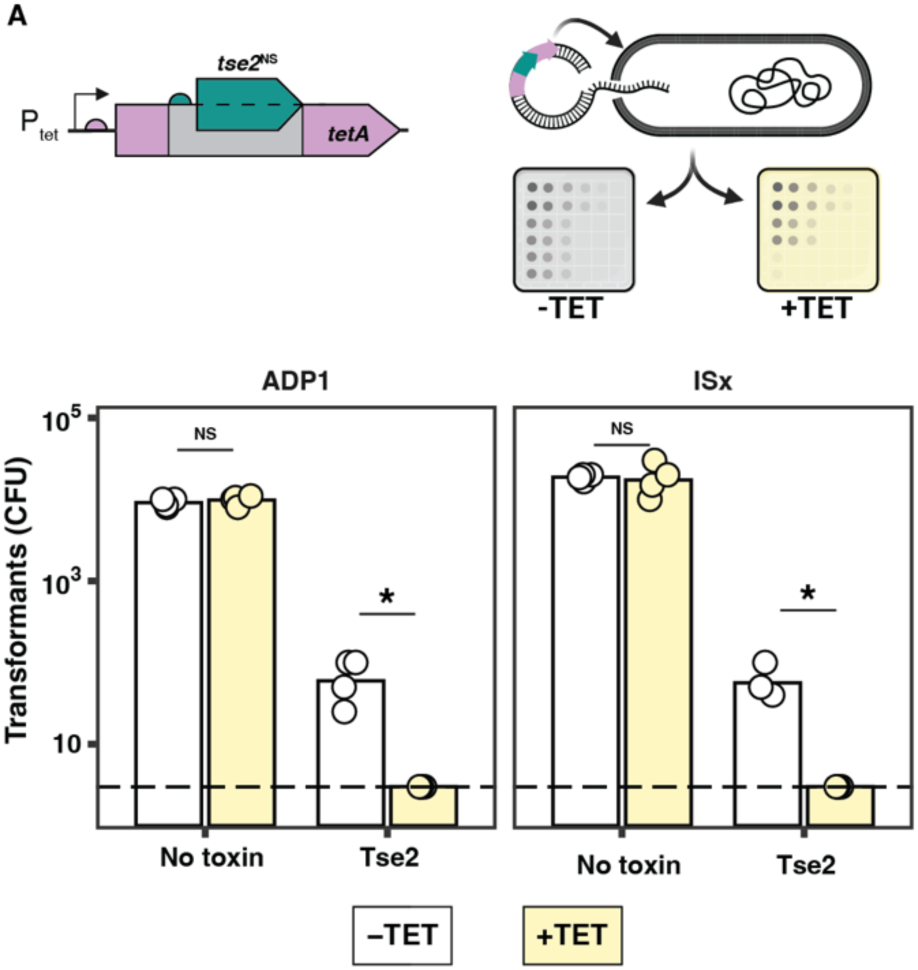
Overlapping toxins limit HGT by natural transformation. (A) Broad-host-range plasmids with *tetA* and with or without the middle *tse2*^NS^ toxin insert were introduced into *A. baylyi* strains ADP1 (left) and ISx (right) by natural transformation. (B) CFUs obtained after overnight incubation in selective conditions. The toxin insert decreased transformants, and inclusion of +TET selection reduced the escape rate below our detection limit. Asterisks indicate statistical differences between groups (ANOVA followed by Tukey HSD, 𝑝 < 0.05)

### Construction of other overlapping gene pairs

To test the extensibility of this approach, we also created additional overlaps of *hicA* and *tse2* within an alternative outer gene, *cmlA*, the chloramphenicol (CHL) resistance gene from Tn*1696* in *Pseudomonas aeruginosa*^34^. As with *tetA*, we selected 3 insertions locations: one N-terminal (“early”, after F39), one in a middle cytoplasmic loop (“middle”, after G200), and one C-terminal (“late”, after R398). We inserted the *hicA*^NS^ and *tse2*^NS^ alleles into these locations and performed assays for toxic function and CHL MIC (**Figure S5A, B**). In contrast to the *tetA* overlapping constructs, in *cmlA* the overlapping toxins functioned only in the early and operon configurations. This correlates well with the *gfp*^NS^ output at each position within *cmlA* (**Figure S5C**), suggesting that varied inner gene expression from different overlap locations controls inner gene function. The CHL MIC for these overlapped constructs followed the trend seen in *tetA*, with early insertions generally being the most disruptive to *cmlA* function, although not when *gfp*^NS^ is the inner gene.

We also created overlapping genes using cytotoxin genes *colE1* (187 aa) and *colE2* (128 aa) as the inner genes within outer genes *tetA* and *cmlA* (**Figure S6-S7**); however, the functionality of the inner (toxin) genes was limited in each case. The recoded, non-overlapped *colE1*^NS^ (requiring 9 synonymous mutations to remove internal stop codons) had nearly abolished toxin activity compared to *colE1*^WT^ based on CFU counts from transformation assays (**Figure S6**).

Interestingly, most colonies from *colE1*^NS^ exhibited a “ghost” phenotype in that they appeared normal sized but did not grow in liquid medium, suggesting some partial function of *colE1*^NS^. In contrast, the recoded, non-overlapped *colE2*^NS^ (requiring 4 synonymous mutations to remove internal stop codons) exhibited similar toxicity to *colE2*^WT^ (**Figure S7**). However, after insertion into *tetA* or *cmlA* at all locations, *colE2*^NS^ showed no toxicity when transformed into susceptible hosts. The cause of this failure in unclear and does not clearly correlate with expected expression level. As with other constructs, the *tetA* and *cmlA* alleles retain partial functionality that is impacted by overlap location. Despite the failure of these additional toxins to create robustly functional overlapping genes, they demonstrate the relative ease with which new overlapping genes can be prototyped and offer an opportunity to understand the failure modes of overlapping genes.

## Methods

### Bacterial culture

For a list of all bacterial strains used in this work see Table S1. *E. coli* NEB 5-alpha cells (Cat# C2987H) were used for routine cloning and assays of non-toxic plasmids. *E. coli* was grown in LB broth (Teknova Cat# L8000) or on agar plates (Teknova Cat# L1000) at 37 °C, shaking when appropriate. For antibiotic selection in *E. coli* we generally used kanamycin (KAN, 25 µg/mL), tetracycline (TET, 20 µg/mL), or chloramphenicol (CHL, 20 µg/mL).

### Plasmid construction

Construction of new plasmids was carried out by NEB HiFi assembly (Cat #E2621S) using Q5 PCR amplified fragments (Cat #M0494S) or IDT synthesized dsDNA fragments (“eBlocks”). Non-toxic plasmid assemblies were transformed into NEB 5-alpha competent cells (Cat # C2987H) according to manufacturer’s instructions. Putatively toxic plasmid assemblies were transformed into the corresponding immune strain, created by constitutively expressing a corresponding antitoxin from the Tn7 *att* site^35^. Immune competent cells were created using Zymo Mix-and-Go (Cat #T3001) and transformed accorded to manufacturer’s instructions. All constructed plasmids were initially verified by sanger sequencing at junctions (Azenta, www.azenta.com) and further verified by whole plasmid sequencing (Plasmidsaurus, www.plasmidsaurus.com).

The progenitor p15A plasmid is pTH40^21^, which we used to create pTH-tetA and pTH-cmlA, which contain (1) wild-type *tetA* or *cmlA* alleles, (2) a kanamycin resistance allele, and (3) low level constitutive gfp fluorescence. Wild-type *tetA* and *cmlA* alleles were synthesized as overlapping eBlocks and inserted into pTH40. Overlapping insert genes and their associated RBS sequence (*hicA*, *colE1*, *colE2*, *tse2*) were ordered as eblocks and inserted into 4 locations of pTH-*tetA* and pTH-*cmlA*. To create the overlapped GFP variants, we first removed GFP from the pTH-*tetA* and pTH-*cmlA* plasmids to create pTH-*tetA*-dGFP and pTH-*cmlA*-dGFP. Then, we ordered recoded GFP as an eblock and inserted it into the same four overlap locations of pTH-*tetA*-dGFP and pTH-*cmlA*-dGFP. To create broad-host-range versions of pTH-*tetA* and overlapped derivatives, we combined PCR fragments of pBTK401 (rsf1010 origin)^30^ with PCR fragments from pTH-*tetA* derivatives that include the *tetA*, *gfp*, and *aphA* (kanamycin resistance). All plasmid sequences are available in Dataset 4.

### Growth and fluorescence assay

Growth and fluorescent output of *E. coli* strains were measured in 96 well flat bottom growth plates (Corning Cat #CLS3631 or #CLS3378) using a BioTek Synergy HTX Multimode Reader plate reader Gen5 v3.11 control and analysis software. We measured GFP at excitation 485/20 nm and emission 528/20 nm with manual gain at 50. Reported GFP values are normalized by OD_600_ measured after blank subtraction.

### Minimum Inhibitory Concentrations (MIC)

MICs were measured using Liofilchem MIC test strips for tetracycline or chloramphenicol (0.016-256 µg/mL, Cat# 921140 and Cat# 920750) according to manufacturer’s instructions. Individual colonies for each strain were grown overnight in 3 mL LB cultures with kanamycin at 37 °C. The next day a sterile cotton swab was used to spread confluent culture onto a pre-dried agar plate. After further drying the plate for 10 minutes, a single MIC strip was placed in the middle of the agar plate and the plate then incubated overnight at 37 °C. MIC was read the next morning after 18 hours of growth by identifying the maximum concentration of antibiotic that allowed for visible growth.

### Transformation toxicity assays

Transformation toxicity assays were performed by transforming purified toxic plasmids into commercial recipient cells and comparing observed transformants to a control transformation with a non-toxic plasmid performed at the same time. Transformations used the NEB recommended protocols for chemical transformation (NEB 5-alpha, Cat# C2987H) or electroporation (NEB 10-beta, Cat# C3019H, 1.8 kV) with 50 ng of plasmid DNA. CFUs were counted after overnight incubation on selective plates, and only regular sized and normal appearing colonies counted.

### qRT-PCR quantification of mRNA expression

*E. coli* strains were grown in 50 mL LB with kanamycin at 37 °C until mid-exponential phase (OD_600_ = 0.25-0.4). 10 mL of bacteria were harvested by centrifugation and lysed using lysozyme and Proteinase K. RNA was isolated from lysates using RNeasy columns (Qiagen). Total RNA was reverse-transcribed using hexameric random primers and SuperScript™ III Reverse Transcriptase (ThermoFisher), according to the manufacturer’s instructions. Real-time quantitative analysis was performed with the LUNA Universal qPCR reagent (New England Biolabs) according to the manufacturer’s instructions, using a CFX Opus Real-Time PCR Systems (BioRad). Two sets of primer pairs were used for detection of *gfp*, *aphA* (kanamycin resistance), and *tetA* regions between the early-middle (“upper”) and middle-late (“lower”) positions. Primers for qRT-PCR were purchased from Integrated DNA Technologies (IDT) and are listed in Table S3. Relative changes were calculated with the efficiency-corrected ΔΔCq method.

### Western blotting

*E. coli* strains were grown in 50 mL LB with kanamycin at 37 °C until mid-exponential phase (OD_600_ = 0.25-0.4). To 1.3 mL of culture, trichloroacetic acid was added to a final of 13% vol/vol, kept on ice overnight, and precipitated by centrifugation. Pellets were washed with ice-cold acetone and resuspended in 1× SDS-PAGE Buffer. Green fluorescent protein (GFP) was detected using a 1:2500 dilution of GFP rabbit antibody (rabbit) Rockland, #600-401-215, while GroEL protein levels were measured using a 1:10,000 dilution of anti-GroEL rabbit antibody (Sigma-Aldrich Cat# G6532). Detection of primary antibodies was performed with a fluorescent secondary antibody (Rabbit IgG (H&L) Antibody DyLight™ 680 Conjugated (Rockland, #611-144-002). Immunoblots were scanned at the appropriate wavelengths for detection. Western blots were quantified using ImageJ software with background subtraction, each band was normalized to GroEL loading control.

### Purification of GFP and variants

To purify the ∼48 kDa GFP variant from NEB 5-alpha expressing *gfp*^NS^ in the middle position of *tetA*, the strain was diluted 250× fold into 1 L of LB with kanamycin and grown shaking at 250 rpm at 37 °C until late exponential phase (OD_600_ = 0.5-0.6). Cells were pelleted at 6,000 × g for 10 min at 4 °C and frozen at -80 °C until further processing. Cells were resuspended in 10 mL of BugBuster HT containing sodium dodecyl sulfate (SDS) at a final concentration of 0.25%, and lysed by nutation at 26 °C for 60 min. Cell debris was pelleted at 7000 × g for 20 min at 4 °C. 250 μL of GFP-Trap Agarose (ChromoTek, #gta-10) was added to the cell supernatant in a separate tube. GFP and GFP variants were bound to the agarose for 60 min at 4 °C, and washed with 10 mL of Buffer A (10 mM Tris-HCl pH 7.5, 150 mM NaCl, 0.5 mM EDTA), followed by 10 mL of Buffer B (10 mM Tris-HCl pH 7.5, 150 mM NaCl, 0.5 mM EDTA, 0.1% SDS, 1% Triton X-100, 1% deoxycholate), followed by 10 mL of Buffer C (10 mM Tris-HCl pH 7.5, 150 mM NaCl, 0.05% Nonidet P40 Substitute, 0.5 mM EDTA). Protein was eluted with 3 x 250 μL of 200 mM glycine pH 2.5. Eluted proteins were concentrated and prepared for mass spectrometry by the chloroform-methanol extraction^36^, separated by SDS-PAGE, and visualized with Coomassie Brilliant Blue. To ensure the ∼48 kDa GFP variant was correctly excised for mass spectrometry analysis, purification of GFP from lysates of NEB 5-alpha expressing *gfp*^NS^ in the operon position was performed in parallel. Protein from five biological replicates were pooled together, and the sequence of the ∼48 kDa GFP variant was analyzed by MS Bioworks LLC (#MSB-01; Ann Arbor, MI). Peptides obtained through mass spectrometry were queried against the amino acid sequence of the *tetA::gfp*^NS^ where *gfp*^NS^ is in the middle position. To identify the region of frameshift, peptide masses were matched to a theoretical molecular weight library generated from a -1 frameshift of each nucleotide in *tetA* starting 7 bp after the ATG start codon of *gfp*^NS^.

### Identifying toxin-inactivating mutations

Individual bacterial colonies were grown to saturation in 3 mL LB, plasmids isolated by miniprep (Zymo Zyppy Plasmid Miniprep, Cat# D4036), and then plasmids sent for long-read nanopore sequencing at Plasmidsaurus (www.plasmidsaurus.com). For results shown in Figure 5, the provided raw reads were processed through a breseq^37,38^ (v0.38.1) pipeline using bowtie2^39^ (v2.5.1) and an appropriate plasmid reference sequence. The NEB 5-alpha genome (accession #CP017100) was annotated for mobile elements using Ises_scan^40^ (v1.7.2.3), and then included as a “junction-only” reference for breseq run in nanopore mode with default parameters. Each sample was analyzed twice, once with the “-t” (“targeted”) flag and once without, and all putative mutations were individually verified by manual inspection of mutation-supporting reads and corrected as needed. For results shown in Figure S4, the inactivating mutations were identified by manual inspection of the assembled plasmid sequence.

### Natural transformation assay for *ADP1* and *ISx*

ADP1 and ISx were transformed using natural competence as previously described^32^. Briefly, frozen stocks of ADP1 and ADP1-ISx were streaked onto LB plates and incubated overnight at 30 °C. Individual colonies were picked into 3 mL LB and incubated overnight at 30 °C with shaking at 200 rpm. 1 mL of each culture was transferred to a 1.5 mL centrifuge tube and centrifuged for 5 min at 3381 × g, culture media removed, and then cell pellets resuspended in 1 mL of fresh LB. 60 μL of this suspension was spotted onto an LB agar plate and combined with 10 μg of Rolling Circle Amplification (RCA) amplified plasmid diluted 1:3 in purified H_2_O. RCA reactions were performed using NEB phi29-XT RCA Kit (Cat# E1603S) according to manufacturer’s instructions with 10 ng purified plasmid as template. Agar plates were incubated at 30 °C overnight and then resuspended in 500 μL of medium and serially diluted for quantification by CFU.

### Statistical analysis and data visualization

Statistical analysis and data visualization were performed in R (v4.4.0, http://www.r-project.org) using tidyverse tools^41^. Statistical analyses of CFU data were generally performed on log-transformed data and statistical tests are specified in figure captions. Portions of Figure 1, Figure 4, and Figure 6 were created using Biorender (https://BioRender.com/w20p944).

### Data and material sharing

Plasmids and strains from this work are available by request. Plasmid sequences are available in Dataset 4. Nanopore sequencing reads used to call mutations shown in Figure 5 are available at Bioproject PRJNA1182389.

## Discussion

In this work, we introduce a simple, effective approach to create new overlapping genes: overlapping, alternate-frame insertion (OAFI). Using OAFI, we created multiple sets of synthetic overlapping genes, identified constructs in which both gene products are functional, and examined how overlap position and selection pressure impact the expression and evolution of both overlapping partners. Given their ease of creation and demonstrated utility, OAFIs represent a general-purpose approach to improve genetic stability of engineered DNA and limit HGT from synthetic microbes into native microbial communities^42^.

OAFI balances generalizability and ease of construction with the mutational robustness of resulting overlapping sequences. Two methods, CAMEOS and RiboSor^13,14^, have been reported for generating synthetic overlapping genes with some experimental validation. CAMEOS designs “full” gene overlaps, in which a smaller gene entirely overlaps within a larger gene. However, this method is computationally intensive and requires extensive screening to identify functional genes. RiboSor creates a partial overlap between a gene and a downstream partner, but this design limits evolutionary coupling between the partners to the overlapping region (e.g., inactivating mutations within a downstream partner gene do not affect the function of the upstream gene). We predict that our OAFI method will be more easily applied to diverse gene pairs without extensive protein redesign, while still maintaining evolutionary coupling between the overlapping genes. One might expect OAFIs to have less mutational robustness than a “full” gene overlap, due to the presumably dispensable nature of the amino acids added in-frame to the outer gene, but direct comparisons of the stability of overlapping genes created through alternate frames have not previously been made. Despite the relative simplicity of OAFI, we found that selection for the outer gene in our middle *tetA::tse2*^NS^ construct reduced toxin escape > 1000- fold. This mutational protection compares favorably to a previously published “full” overlapping gene pair *ilvA::relE* created by CAMEOS^11^, in which selection for the outer gene (*ilvA* function) reduced toxin escape by approximately 10-fold. Further, our results show that the OAFI method protects against deletions mediated by homologous recombination and insertions of mobile elements, two relatively frequent mutations observed in *E. coli*^43–45^, although we did not observe protection against SNPs, as observed with the previous *ilvA::relE* pair. Thus, we suspect the protection afforded by OAFI could usefully enhance the stability of diverse engineered constructs despite not protecting from SNPs.

We do not explicitly design the inserted sequence in the original frame of the outer gene, which makes OAFI intrinsically flexible. This flexibility makes the approach immediately extensible to comprehensive studies designed to elucidate the epistatic effects of overlapping reading frames on gene expression. While natural gene overlaps have been studied to understand their impact on translation in viruses^46^, there are too few such examples to rigorously dissect the impact of gene overlap on protein translation. OAFI enables a direct, high-resolution assessment of how specific sequence characteristics (e.g., internal and external RBS strength, rare codon usage, ribosome slippage sites, secondary mRNA structure, internal ORF position) impact the translation of both genes in diverse overlapping pairs. Studies have shown that mRNAs have intrinsic ORF-wide secondary structure that strongly influences translation efficiency, and that the immediate vicinity of the start sites is relatively unstructured for most ORFs^47^. This suggests that RNA secondary structure and positional effects can significantly impact the translation of overlapping genes. Indeed, we observed that inner gene translation varies dramatically depending on its position within the outer gene. Furthermore, in at least one design (*gfp*^NS^ in the middle position of *tetA*), we found evidence of translational frame-shift, a potential consequence of ribosome stalling and/or collision^48,49^. We anticipate that recent methods for saturated insertional mutagenesis^50,51^ could be combined with OAFI to systematically evaluate the positional, structural, and sequence-specific impacts on the translational mechanics of overlapping genes.

Beyond translation mechanics, OAFI constructs provide a means to explore the evolution of overlapping genes and their role in modulating the evolutionary fate of one or both partner genes. Even though the inner gene is translated into a presumably non-functional amino acid sequence in the outer ORF, we found that selection for the outer gene nevertheless influences the evolutionary fate of the inserted sequence. Specifically, overlapping pairs with conditionally essential outer genes and a cytotoxic internal gene restricted the most common types of inactivating mutations in the internal gene, namely large deletions and mobile element insertions. Replacing the internal gene with an essential (or conditionally essential) gene would allow for selection of both genes simultaneously and may reveal unexpected evolutionary novelties that enhance the function of both overlapping genes. Whether essentiality of the inner gene can also be harnessed to increase genetic stability—that is, alter the mutational trajectory of a costly outer gene—remains to be investigated. Ultimately, the novel evolutionary paths of OAFI pairs could mimic those of fully overlapped viral genes, wherein mutation rates appear to be fine-tuned by overlapping essential regions of one partner with mutable regions of the other^52^. OAFI designs thus provide an engineerable model and a fruitful testbed to understand the evolutionary dynamics of overlapping genes under distinct evolutionary pressures, particularly in cases where both genes are under strong positive or purifying selection.

While inactivation of a costly internal gene is inevitable over long timescales, we anticipate that overlapping insertions can prolong genetic stability to meet desired outcomes within applications in agriculture and industrial microbiology, among others^53^. Moreover, applying directed evolution or similar approaches to overlapping genes has the potential to optimize the function of both genes to improve genetic circuit lifespan^20^. We took no effort to optimize the alternate frame amino acids of the inner genes, but it is likely that performance of the outer gene could be enhanced by intentional design of the added amino acids to reduce unstructured regions or encode specific secondary or tertiary constraints. Our initial results also demonstrate that overlapping gene pairs can prevent the transfer of antibiotic resistance genes among multiple bacterial species. Given this success, we envision that our method could be readily implemented as a biocontainment strategy to prevent the spread of antibiotic resistance genes via HGT from various biotechnological activities. Our results, however, also highlight the need to screen multiple toxins and configurations when designing these systems to ensure functionality and mutational robustness in relevant bacteria^21^. OAFIs thus have the potential to provide the benefits of natural overlapping genes, translating synthetic biology advances to real-world applications.

## Acknowledgements

We thank Jeffrey Barrick for providing strains ADP1 and ISx. Funding was provided by U.S. Department of Energy (DOE), Office of Science, Office of Biological and Environmental Research, Secure Biosystems Design Program, Lawrence Livermore National Laboratory’s BioSecure Scientific Focus Area #SCW1710. All work carried out at Lawrence Livermore National Lab under the auspices of the U.S. Department of Energy at Lawrence Livermore National Laboratory under Contract DE-AC52-07NA27344 (LLNL-JRNL-2000896).

## Supplemental

Table ST1: Strains

Table ST2: Plasmids

Table ST3: Primers

Dataset 1: Scaffold (.sf3) results of MS/MS

Dataset 2: List of mutations identified by sequencing, Tet_10_ selection

Dataset 3: List of mutations identified by sequencing, Tet_5_ selection

Dataset 4: Plasmid sequences

Figure S1: Relative fluorescence of GFP in *tetA*::gfp^NS^ strains.

Figure S2: mRNA abundance of overlapping genes *tetA*::*gfp*^NS^

Figure S3: The junction of a TetA-GFP fusion identified through mass spectrometry.

Figure S4: *Tse2*^NS^ inactivation under Tet_5_ selection.

Figure S5: Function of *tse2*^NS^, *hicA*^NS^, *gfp*^NS^ inserts in *cmlA*.

Figure S6: Function of *colE1* toxin inserted into *tetA* and *cmlA*.

Figure S7: Function of *colE2* toxin inserted into *tetA* and *cmlA*.

