## Supplemental information for "Creating overlapping genes by alternate-frame insertion"

**This file includes:**

Figures S1 to S7
Tables S1 to S3

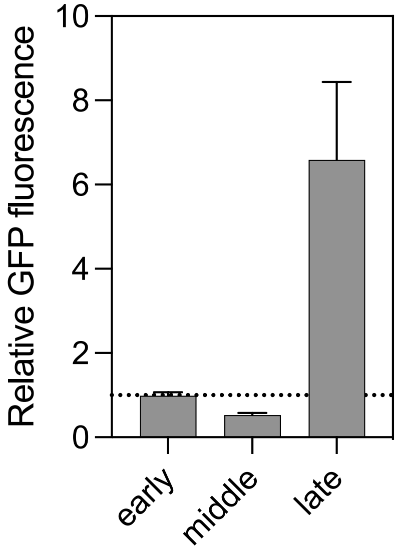


**Figure S1. Relative fluorescence of GFP in *tetA*::*gfp*[NS] strains**. Quantification of immunoblot analysis from Figure 4B. Error bars indicate standard deviation.

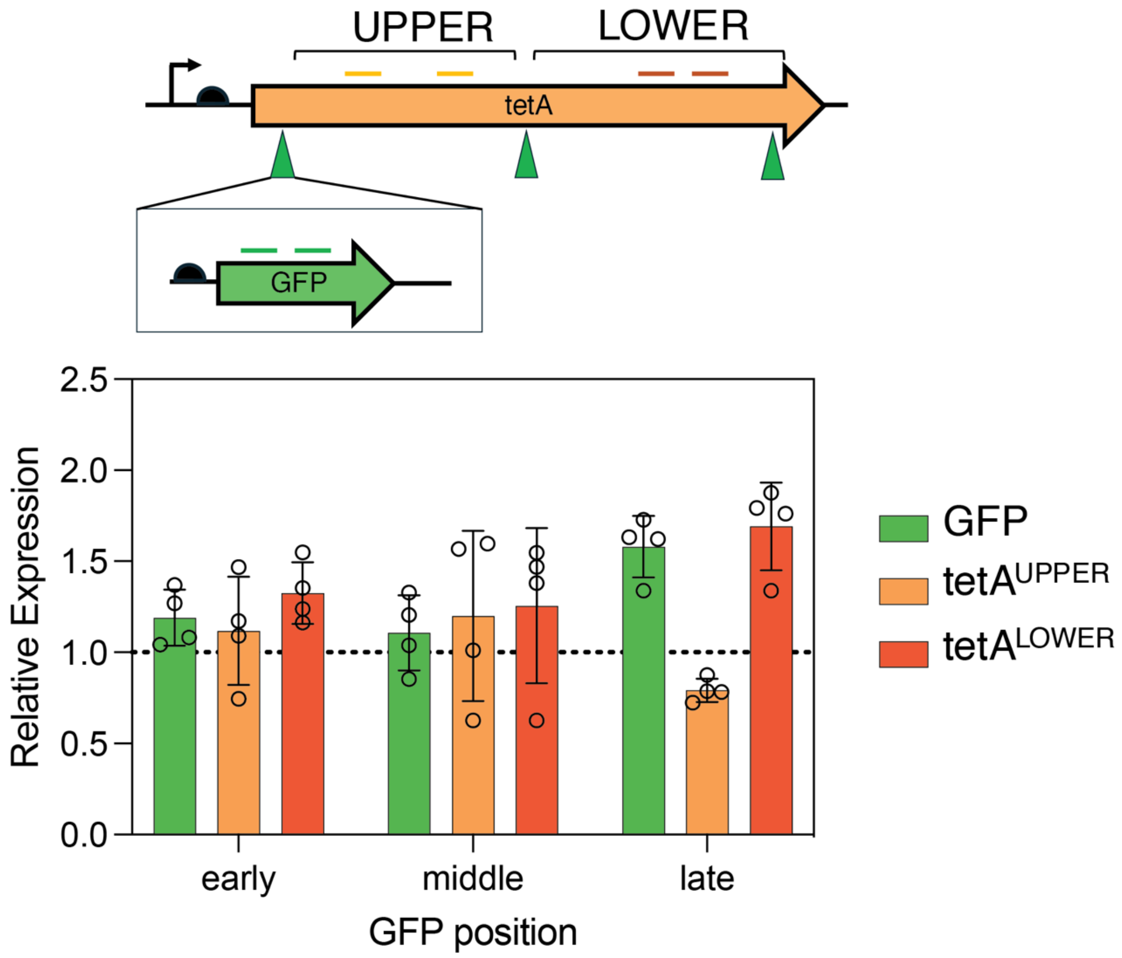


**Figure S2. mRNA abundance of overlapping genes *tetA*::*gfp*$^{NS}$.** Dotted line is normalized expression of the wild-type *tetA* allele. No differences are statistically significant ($p > 0.05$). Error bars indicate standard deviation.

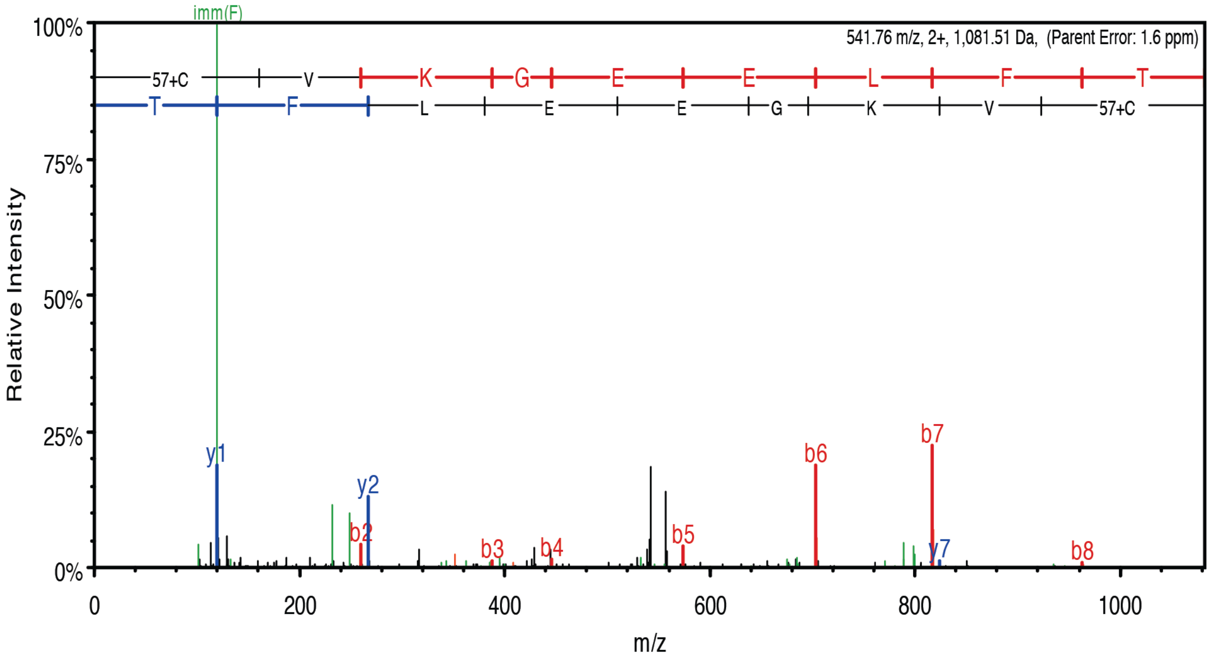


**Figure S3. The junction of a TetA-GFP fusion identified through mass spectrometry.** The MS/MS spectrum corresponding to a frameshift of the TetA-GFP$^{mid}$ construct between amino acid positions 201 and 202 of purified protein is shown.

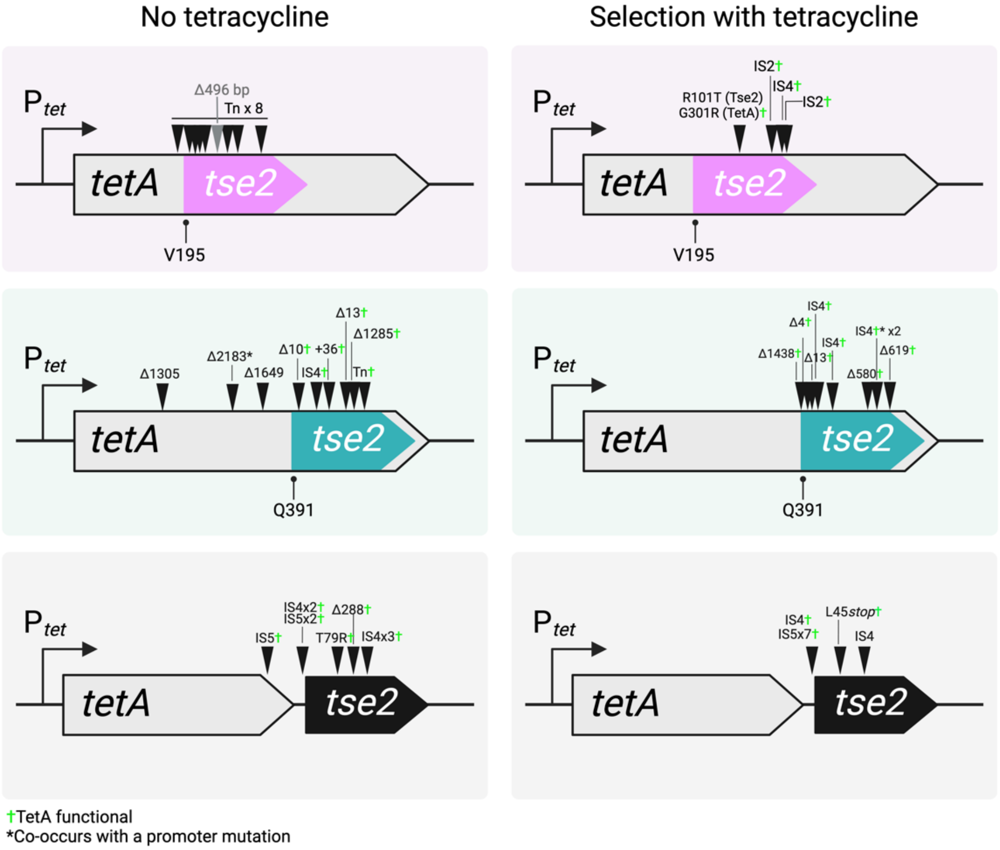


**Figure S4. Tse2$^{NS}$ inactivation under Tet$_5$ selection.** For all mutations see Dataset 3.

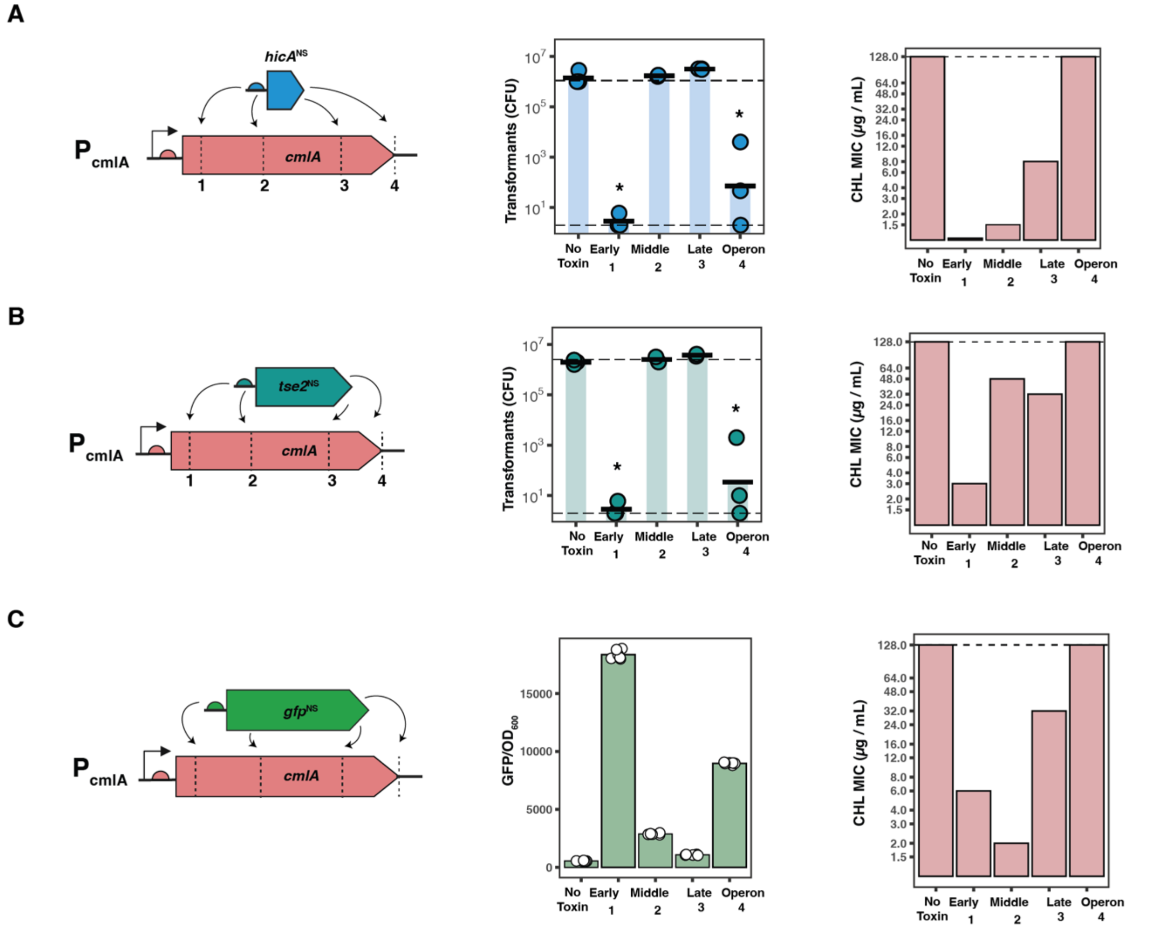


**Figure S5. Function of *tse2*$^{NS}$, *hicA*$^{NS}$, *gfp*$^{NS}$ inserts in *cmlA*.** (A) Transformation assays (middle) and CHL MIC (right) of overlapping constructs with *hicA*$^{NS}$ inserted into *cmlA*. HicA inserts are toxic only in the early and operon configurations. (B) Transformation assays (middle) and CHL MIC (right) of overlapping constructs with *tse2*$^{NS}$ inserted into *cmlA*. Tse2 inserts are toxic only in the early and operon configurations. (C) Fluorescence (middle) and CHL MIC (right) of overlapping constructs with *gfp*$^{NS}$ inserted into *cmlA*. For transformation assays upper dotted line corresponds to number of transformants obtained with plasmid bearing wild-type CmlA with no toxin. Lower dotted line corresponds to approximate limit of detection. For MIC assays dotted line corresponds to MIC of wild-type CmlA allele. Asterisk indicates statistically significant differences between groups (ANOVA followed by Tukey HSD, $p < 0.05$), except in fluorescence assay where all groups are significantly different from one another.

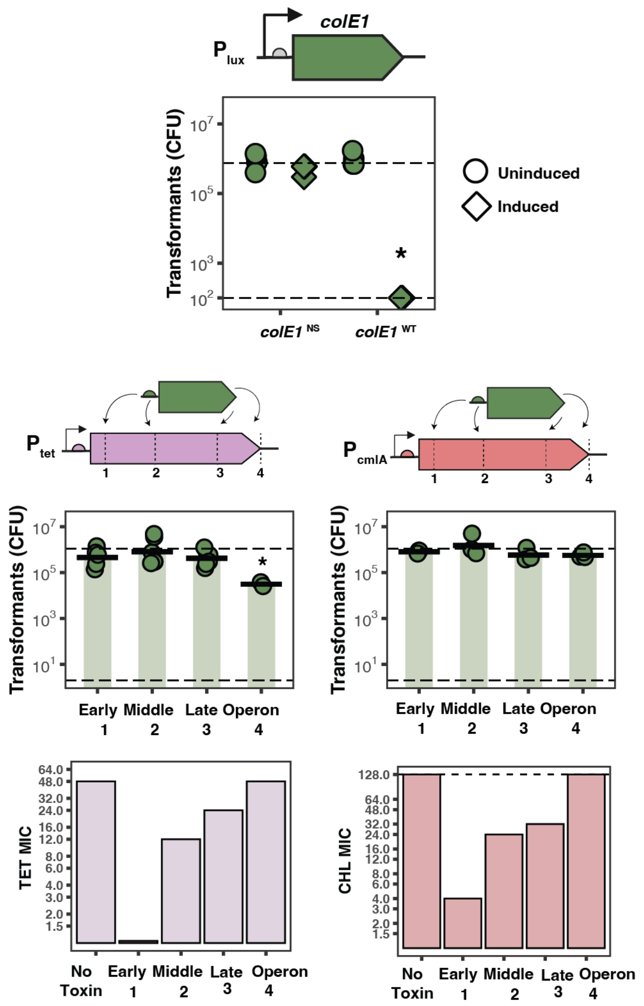


**Figure S6: Function of *colE1* toxin inserted into *tetA* and *cmlA*.** Transformation toxicity and MIC assays for *colE1* toxin inserted into *tetA* and *cmlA*. Experimentation and data analysis and presentation identical to Figure 2 and Figure S5.

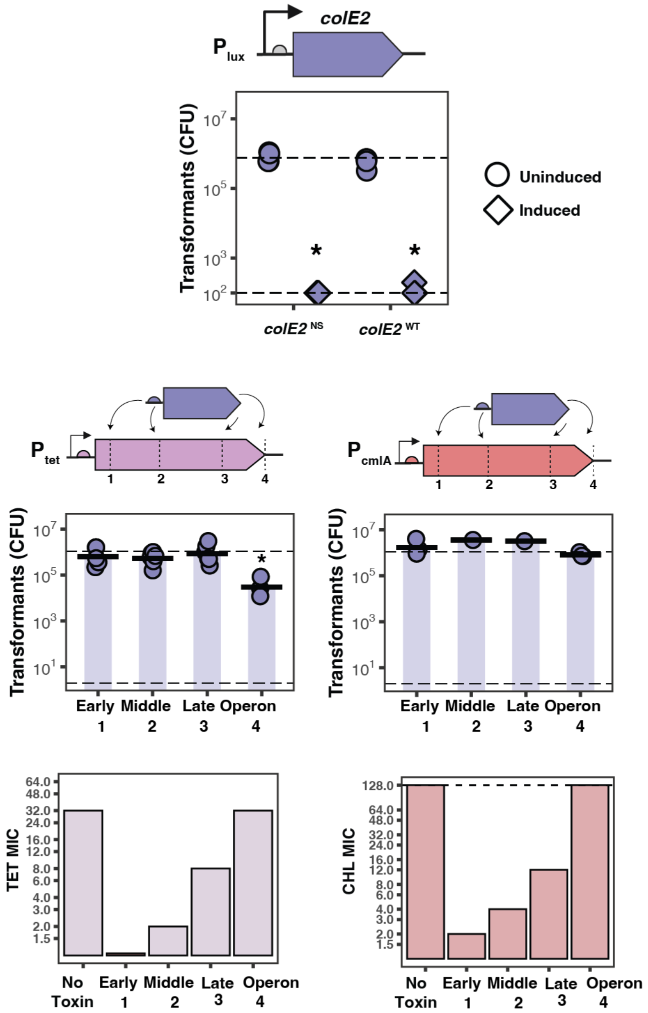


**Figure S7: Function of *colE2* toxin inserted into *tetA* and *cmlA*.** Transformation toxicity and MIC assays for *colE2* toxin inserted into *tetA* and *cmlA*. Experimentation and data analysis and presentation identical to Figure 2 and Figure S5.

**Table S1**. Bacterial strains

| Species and strain | Source | ID |
|---|---|---|
| *Escherichia coli* NEB 5-alpha | New England Biolabs | CAT# C2987H |
| *E. coli* NEB 10-beta | New England Biolabs | CAT# C3019H |
| *Acinetobacter baylyi* ADP1 | [1] | ATCC#33305 |
| *A. baylyi* ADP1-ISx | [2] | N/A |
| *E. coli* NEB 5-alpha *attTn7::CP25-hicB* | This study | N/A |
| *E. coli* NEB 5-alpha *attTn7::CP25-tsi2* | This study | N/A |
| *E. coli* NEB 5-alpha *attTn7::CP25-immE1* | This study | N/A |
| *E. coli* NEB 5-alpha *attTn7::CP25-immE2* | This study | N/A |

**Table S2.** Plasmids.

| Name | Description | Source |
|---|---|---|
| pTH_tetA | Wild-type *tetA* on p15a origin with kanamycin resistance allele | This study |
| pTH_cmlA | Wild-type *cmlA* on p15a origin with kanamycin resistance allele | This study |
| pBLxG2_hicA | $hicA^{WT}$ under PLuxB induction on pbbr1 origin with kanamycin resistance | This study |
| pBLxG2_hicA_NS | $hicA^{NS}$ under PLuxB induction on pbbr1 origin with kanamycin resistance | This study |
| pBLxG2_tse2 | $tse2^{WT}$ under PLuxB induction on pbbr1 origin with kanamycin resistance | This study |
| pBLxG2_tse2_NS | $tse2^{NS}$ under PLuxB induction on pbbr1 origin with kanamycin resistance | This study |
| pBLxG2_colE1 | $colE1^{WT}$ under PLuxB induction on pbbr1 origin with kanamycin resistance | This study |
| pBLxG2_colE1_NS | $colE1^{NS}$ under PLuxB induction on pbbr1 origin with kanamycin resistance | This study |
| pBLxG2_colE2 | $colE2^{WT}$ under PLuxB induction on pbbr1 origin with kanamycin resistance | This study |
| pBLxG2_colE2_NS | $colE2^{NS}$ under PLuxB induction on pbbr1 origin with kanamycin resistance | This study |
| pTH_tetA_S36_hicA | *tetA*::$hicA^{NS}$ early | This study |
| pTH_tetA_V195_hicA | *tetA*::$hicA^{NS}$ middle | This study |
| pTH_tetA_Q391_hicA | *tetA*::$hicA^{NS}$ late | This study |
| pTH_tetA_operon_hicA | *tetA*::$hicA^{NS}$ operon | This study |
| pTH_tetA_S36_tse2 | *tetA*::$tse2^{NS}$ early | This study |
| pTH_tetA_V195_tse2 | *tetA*::$tse2^{NS}$ middle | This study |
| pTH_tetA_Q391_tse2 | *tetA*::$tse2^{NS}$ late | This study |
| pTH_tetA_operon_tse2 | *tetA*::$tse2^{NS}$ operon | This study |
| pTH_tetA_S36_gfp | *tetA*::$gfp^{NS}$ early | This study |
| pTH_tetA_V195_gfp | *tetA*::$gfp^{NS}$ middle | This study |
| pTH_tetA_Q391_gfp | *tetA*::$gfp^{NS}$ late | This study |
| pTH_tetA_operon_gfp | *tetA*::$gfp^{NS}$ operon | This study |
| pTH_tetA_rsf1010 | Wild-type *tetA* on rsf1010 origin with kanamycin resistance allele | This study |
| pTH_tetA_V195_tse2_rsf1010 | *tetA*::$tse2^{NS}$ middle on rsf1010 origin with kanamycin resistance allele | This study |
| pTH_40 | Source of p15a origin backbone | [3] |
| pBLxG2 | Source of pbbr1 origin with PLuxB induction backbone | [4] |
| pBTK403 | Source of rsf1010 origin backbone | [5] |

**Table S2.** Plasmids. (cont)

| Name | Description | Source |
|---|---|---|
| pTH_cmlA_F39_hicA | *cmlA*::hicA$^{NS}$ early | This study |
| pTH_cmlA_G200_hicA | *cmlA*::hicA$^{NS}$ middle | This study |
| pTH_cmlA_R398_hicA | *cmlA*::hicA$^{NS}$ late | This study |
| pTH_cmlA_operon_hicA | *cmlA*::hicA$^{NS}$ operon | This study |
| pTH_cmlA_F39_tse2 | *cmlA*::tse2$^{NS}$ early | This study |
| pTH_cmlA_G200_tse2 | *cmlA*::tse2$^{NS}$ middle | This study |
| pTH_cmlA_R398_tse2 | *cmlA*::tse2$^{NS}$ late | This study |
| pTH_cmlA_operon_tse2 | *cmlA*::tse2$^{NS}$ operon | This study |
| pTH_cmlA_F39_gfp | *cmlA*::gfp$^{NS}$ early | This study |
| pTH_cmlA_G200_gfp | *cmlA*::gfp$^{NS}$ middle | This study |
| pTH_cmlA_R398_gfp | *cmlA*::gfp$^{NS}$ late | This study |
| pTH_cmlA_operon_gfp | *cmlA*::gfp$^{NS}$ operon | This study |
| pTH_tetA_S36_colE1 | *tetA*::colE1$^{NS}$ early | This study |
| pTH_tetA_V195_colE1 | *tetA*::colE1$^{NS}$ middle | This study |
| pTH_tetA_Q391_colE1 | *tetA*::colE1$^{NS}$ late | This study |
| pTH_tetA_operon_colE1 | *tetA*::colE1$^{NS}$ operon | This study |
| pTH_tetA_S36_colE2 | *tetA*::colE2$^{NS}$ early | This study |
| pTH_tetA_V195_colE2 | *tetA*::colE2$^{NS}$ middle | This study |
| pTH_tetA_Q391_colE2 | *tetA*::colE2$^{NS}$ late | This study |
| pTH_tetA_operon_colE2 | *tetA*::colE2$^{NS}$ operon | This study |
| pTH_cmlA_F39_colE1 | *cmlA*::colE1$^{NS}$ early | This study |
| pTH_cmlA_G200_colE1 | *cmlA*::colE1$^{NS}$ middle | This study |
| pTH_cmlA_R398_colE1 | *cmlA*::colE1$^{NS}$ late | This study |
| pTH_cmlA_operon_colE1 | *cmlA*::colE1$^{NS}$ operon | This study |
| pTH_cmlA_F39_colE2 | *cmlA*::colE2$^{NS}$ early | This study |
| pTH_cmlA_G200_colE2 | *cmlA*::colE2$^{NS}$ middle | This study |
| pTH_cmlA_R398_colE2 | *cmlA*::colE2$^{NS}$ late | This study |
| pTH_cmlA_operon_colE2 | *cmlA*::colE2$^{NS}$ operon | This study |

**Table S3.** Primers

| ID | Use | Source | Sequence (5' to 3') |
|---|---|---|---|
| GFP2_for | qPCR GFP set 1 forward | This Study | TGGTGATGTCAACGGTCATAAG |
| GFP2_rev | qPCR GFP set 1 reverse | This Study | GCAAAGCACTGAACACCATAAG |
| GFP4_for | qPCR GFP set 2 forward | This Study | TGCAGGAACGCACGATTT |
| GFP4_rev | qPCR GFP set 2 reverse | This Study | CAGCTTATGGCCCAGGATATT |
| tetAU3_for | qPCR tetA upper set 1 forward | This Study | GATACCACCTCAGCTTCTCAAC |
| tetAU3_rev | qPCR tetA upper set 1 reverse | This Study | CAATATTTAGCAACGCAGCGATAA |
| tetAU4_for | qPCR tetA upper set 2 forward | This Study | TTGCTCCTTGGCTTGGAAA |
| tetAU4_rev | qPCR tetA upper set 2 reverse | This Study | CTGAAAGCAAACGGCCTAAATAC |
| tetAL3_for | qPCR tetA lower set 1 forward | This Study | CAGGGAGTGATGTCTATCCAAAC |
| tetAL3_rev | qPCR tetA lower set 1 reverse | This Study | TAATCCAAATCCAGCCATCCC |
| tetAL4_for | qPCR tetA lower set 2 forward | This Study | CGCAATTGATAGGCCAAATTCC |
| tetAL4_rev | qPCR tetA lower set 2 reverse | This Study | GGCTATTCTTCCTGCCACAA |
| aphA2_for | qPCR aphA set 1 forward | This Study | CGAACGATGTGACCGATGAA |
| aphA2_rev | qPCR aphA set 1 reverse | This Study | GGATATTCTTCCAGCACCTGAA |
| aphA5_for | qPCR aphA set 2 forward | This Study | TTTCTGCGTCGTCTGCATAG |
| aphA5_rev | qPCR aphA set 2 reverse | This Study | CCAGCCGTTACGTTCATCAT |
